# Affinity Scores: An Individual-centric Fingerprinting Framework for Neuropsychiatric Disorders

**DOI:** 10.1101/2021.11.15.468749

**Authors:** Cassandra M J Wannan, Christos Pantelis, Antonia Merritt, Bruce Tonge, Warda T Syeda

## Abstract

**Background:** Population-centric frameworks of biomarker identification for psychiatric disorders focus primarily on comparing averages between groups and assume that diagnostic groups are (1) mutually-exclusive, and (2) homogeneous. There is a paucity of individual-centric approaches capable of identifying individual-specific ‘*fingerprints*’ across multiple domains. To address this, we propose a novel framework, combining a range of biopsychosocial markers, including brain structure, cognition, and clinical markers, into higher-level ‘*fingerprints*’, capable of capturing intra-illness heterogeneity and inter-illness overlap.

**Methods:** A multivariate framework was implemented to identify individualised patterns of brain structure, cognition and clinical markers based on affinity to other participants in the database. First, individual-level affinity scores defined each participant’s “neighbourhood” across each measure based on variable-specific hop sizes. Next, diagnostic verification and classification algorithms were implemented based on multivariate affinity score profiles. To perform affinity-based classification, data were divided into training and test samples, and 5-fold nested cross-validation was performed on the training data. Affinity-based classification was compared to weighted K-nearest neighbours (KNN) classification. K-means clustering was used to create clusters based on multivariate affinity score profiles. The framework was applied to the Australian Schizophrenia Research Bank (ASRB) dataset.

**Results:** Individualised affinity scores provided a ‘*fingerprint*’ of brain structure, cognition, and clinical markers, which described the affinity of an individual to the representative groups in the dataset Diagnostic verification capability was moderate to high depending on the choice of multivariate affinity metric. Affinity score-based classification achieved a high degree of accuracy in the training, nested cross-validation and prediction steps, and outperformed KNN classification in the training and test datasets.

**Conclusion:** Affinity scores demonstrate utility in two keys ways: (1) Early and accurate diagnosis of neuropsychiatric disorders, whereby an individual can be grouped within a diagnostic category/ies that best matches their fingerprint, and (2) identification of biopsychosocial factors that most strongly characterise individuals/disorders, and which may be most amenable to intervention.

## Introduction

Population-centric frameworks of biomarker identification focus primarily on comparing averages between groups and assume that diagnostic groups are (1) independent, and (2) homogeneous. However, there is a considerable overlap in symptoms across psychiatric disorders, and variability within the spectrum of symptoms of specific disorders^1–3^. Furthermore, there is now increasing understanding of intra-illness heterogeneity, in which individuals who share a diagnosis can present with a varying array of symptoms and disabilities across a range of clinical, cognitive, adaptive behaviour, and social domains^4,5^. Intra-illness heterogeneity, combined with overlap of symptoms and frequent comorbidity of psychiatric disorders, can lead to clinically significant delay in receiving the most appropriate treatment, resulting in increased social and functional disability, family burden and accruing societal economic burden^6^. Consequently, in recent years, there has been an increased focus on the identification of biomarkers that may characterise specific disorders, and aid in diagnosis, subgrouping, and treatment planning.

Various approaches have been developed to identify hierarchical biomarkers using individual base-level measures, such as regional brain volumes, stress measures and cognitive variables. Such approaches can be broadly classified into three categories: 1) multivariate pattern identification, 2) clustering analyses for machine learning and prediction, and 3) normative models. Although these approaches have contributed strongly towards increasing our understanding of biopsychosocial underpinnings of psychiatric disorders, they offer limited clinical translation. Group-level analyses (e.g., pattern identification, clustering) fail to capture individual complexity spanning over multiple clinical domains, whereas normative approaches are typically performed within a single domain of interest and are based on the assumption that there is a true healthy population to act as a reference group.

Recent approaches have attempted to address some of these limitations by taking a more individual-centric approach to disease identification and classification. One method, developed for use in Alzheimer’s disease, is the Disease State Index (DSI)^7^. The DSI is a statistical modeling and data visualisation system that computes an estimate of an individual’s Alzheimer’s disease state by comparing their biomarker data to previously diagnosed cases. This approach enables individualised multivariate quantification of the disease state; however, an important limitation is that scores are referenced to multiple target populations whilst incorporating information from the entire distributions, and are, therefore, less sensitive to the localised information most closely relevant to the individual-of-interest^8^. One method sensitive to localised information in the data with respect to the participant-of-interest is K-Nearest Neighbours (KNN), a supervised machine learning algorithm^9,10^. The KNN algorithm assumes that similar data points (e.g., diagnoses) are close to each other, and therefore classifies new data points based on their proximity to *K* nearest neighbours. This approach offers limited reliability in the presence of outliers, as their nearest neighbours could be situated far from them. The weighted KNN algorithm attempts to minimize this effect by applying inverse distance weights to the neighbours. Conversely, in densely populated neighbourhoods, selecting only first *K* neighbours leads to exclusion of information from other nearby neighbors, which can lead to suboptimal performance, especially when there is large information overlap between diagnostic groups.

In order to address the limitations of current approaches, we propose a novel framework capable of capturing intra-illness heterogeneity and inter-illness overlap through combining a range of biopsychosocial markers into higher-level biopsychosocial ‘*fingerprints*’. In this framework, each individual is scored based on their biopsychosocial marker affinity to different diagnostic groups within a fixed sized neighbourhood, thereby generating overlapping indices of group membership. In the current study, ‘*affinity scores*’ represent biopsychosocial patterns of treatment-responsive and treatment-resistant schizophrenia, whereby we capture each individual’s affinity to multiple diagnostic groups across a wide range of variables. This individual-level approach will have translational value as it allows clinicians to acknowledge comorbidity, indicate severity, disability and prognosis, and tailor intervention strategies to each person’s clinical and biopsychosocial ‘*fingerprint*’. Additionally, we integrate our affinity score framework into two distinct diagnostic verification and classification/prediction algorithms, which can be used as a clinical decision support system designed to enable clinicians to establish a diagnosis and determine a likely prognosis. Our framework will also be compared with an existing framework, K-Nearest Neighbours (KNN) classification, in order to determine the prediction accuracy of our affinity-based classification. Finally, k-means clustering will be applied to multivariate Affinity Score profiles in order to assess the separability of treatment-resistant individuals from healthy controls and individuals with chronic schizophrenia.

## Methods

### Participants

Data from 186 individuals with chronic schizophrenia (age: 37.99 ± 09.98, 69.9% males) and 168 healthy controls (age: 39.74 ± 13.97, 48.2% males) were obtained from the Australian Schizophrenia Research Banks (ASRB), a register of research data collected between 2007-2011 by scientific collaborators across five Australian sites^11^. For the purpose of the current study, participants receiving treatment with clozapine (n = 37, age: 36.62 ± 09.86, 81.1% males) were designated as having treatment-resistant schizophrenia (TRS)^12–14^. ASRB exclusion criteria for participants included: i) a history of organic brain disorder; ii) electroconvulsive therapy in the previous 6 months; iii) current substance dependence; iv) movement disorders; or v) brain injury with post-traumatic amnesia. Healthy controls with a personal or family history of psychosis or bipolar I disorder were also excluded. Detailed information regarding the consent procedures have been published previously^11^. The Diagnostic Interview for Psychosis (DIP)^15^ was used to obtain clinical symptom ratings and confirm diagnoses according to ICD-10 or DSM-IV criteria. Study procedures were approved by the Melbourne Health Human Research Ethics Committee and written informed consent was obtained from all participants.

Participants with at least one missing variable were excluded from the study, resulting in a total sample size of 182 individuals with chronic schizophrenia (Scz), 136 healthy controls (HCs) and 34 TRS individuals.

### Clinical and cognitive measures

The Schizotypal Personality Questionnaire (SPQ)^16^ was used to measure schizotypal symptoms in all participants. The Global Assessment of Functioning (GAF)^17^ was administered to assess current functioning.

Current IQ was measured using the Vocabulary and Matrix Reasoning subtests of the Wechsler Abbreviated Scale of Intelligence (WASI)^18^. Premorbid IQ was measured using the Wechsler Test of Adult Reading (WTAR)^19^. Additional cognitive tasks included Letter Number Sequencing (LNS)^20^, the Controlled Oral Word Association Test (COWAT)^21^, and five subscales (immediate memory, delayed memory, construction, language, attention) from the Repeatable Battery for Assessment of Neuropsychological Status (RBANS)^22^.

### MRI acquisition and processing

T1-weighted (MPRAGE) structural scans were acquired using Siemens Avanto 1.5T scanners located at five different sites in Australia. The same acquisition sequence was used at each site and a Siemens phantom was periodically imaged at each site to evaluate potential inter-site differences. T1-weighted images comprised 176 sagittal slices of 1 mm thickness without gap; field of view = 250 × 250 mm^2^; repetition time = 1980 ms, echo time = 4.3 ms; data matrix size = 256 × 256; voxel dimensions = 1.0 × 1.0 × 1.0 mm^3^.

To estimate cortical thickness, volume, and surface area, images were processed with the FreeSurfer software package version 5.1 (https://surfer.nmr.mgh.harvard.edu)^23^. In brief, preprocessing included intensity normalization, removal of non-brain tissue, transformation to Talairach-like space, segmentation of gray-white matter tissue, and tessellation and smoothing of the white matter boundary. White matter surfaces were then deformed toward the gray matter boundary at each vertex. The entire cortex of each study participant was visually inspected, and inaccuracies in segmentation were manually edited. The cortical surface was then parcellated into 68 regions based on the Desikan-Killiany atlas^24^. Subcortical volumes (thalamus, amygdala, and hippocampus) and estimated total intracranial volume (ICV) were also obtained using the Freesurfer ‘aseg’ parcellation.

### Variable-wise affinity scores

The variable-wise affinity scores are calculated on the combined sample from all diagnostic groups. Each variable is converted into z-scores using mean and standard deviation from the pooled data. The variable-specific affinity scores are calculated using the following algorithm.

1. A variable-specific hop-size, *h_v_*, is determined in the standard deviation units as,

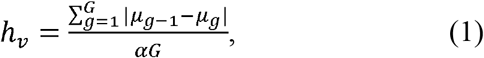

where, *μ_g_* is the mean z-score of group *g, g* = 1, …, *G*, and *G* is the number of diagnostic groups in the study. *α* is a scaling constant.
2. For each participant, *p*, the distance from every other participant in the dataset is calculated in the standard deviation units, and a neighbourhood, 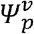, of size 2*h_v_* is defined centered at the z-score of the participant *p*, (*z_p_* ± *h_v_*). Participants within the neighbourhood 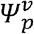 contribute towards the group-wise affinity score, 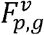, calculated as,

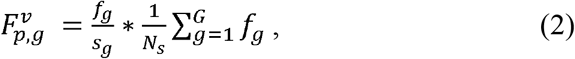

where, *f_g_* is the number of participants from group *g* within 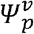, *g* = 1, …, *G, s_g_* is the group sample size and *N_s_* is the total number of participants in the data, 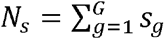. The affinity scores can be used for diagnostic verification (participant *p* is included in 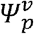) or classification (participant *p* is excluded from 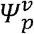).
3. A variable-specific adjacency matrix, ***A**^v^*, is defined as:

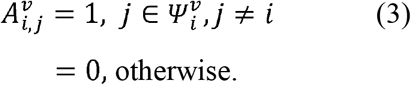 ***A**^v^* is a matrix of size *N_s_* × *N_s_* and *i, j* = 1, …, *N_s_*.

### Multivariate affinity

Multiple metrics can be calculated to assess the multivariate distribution of a participant’s affinity to each diagnostic group in the data.

### Composite multivariate affinity

For each participant, a group-specific composite affinity score is calculated by averaging the affinity scores across all variables,

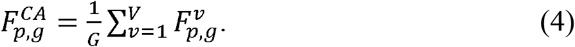

The composite affinity scores quantify the average affinity of *p*th participant to each diagnostic group, *g*, in the data. *C* is the number of diagnostic groups in the data and *V* is the total number of variables.

### Rank-based multivariate affinity

For each variable and participant, affinity scores can be ranked based on the affinity strength to each diagnostic group, such that the group with least affinity score is ranked the lowest. The group-wise rank-based average affinity score is calculated as:

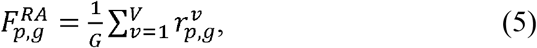

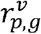 is the variable-wise group rank for *p*th participant and takes values in the range 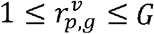.

### Vote-based multivariate affinity

The vote-based affinity score for the *p*th participant is based on a voting system such that each variable votes 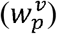 towards the diagnostic group with maximum affinity:

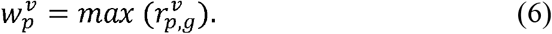

The multivariate vote-based affinity score is calculated using the vote-count, such that the diagnostic group with the most votes scores the highest.

### Common neighbourhood-based affinity

The multivariate common neighbourhood is defined as the number of variables in which a pair of participants share a common neighbourhood. The multivariate common neighbourhood adjacency matrix of size *N_s_* × *N_s_* is defined as

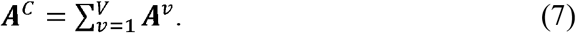

The group-wise common neighbourhood-based affinity for the *p*th participant is calculated as:

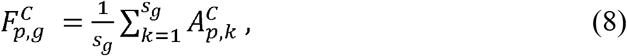

where *g* = 1, …, *G*.

### Common community-based affinity

The adjacency matrix of each variable ***A**^v^* can be used to identify within-variable communities using a clustering algorithm, such as graph clustering with modularity maximization described in (Blondel et al, 2008)^25^. Each participant is assigned a community, and a participant cannot be a member of multiple communities. The participants within the community share maximum affinity to each other, and each community shares least affinity to other communities. The common community-based affinity scores are calculated by the following algorithm:

1. Within variable communities are identified in the adjacency matrix ***A**^v^* using a graph clustering algorithm, such as (Blondel et al, 2008)^25^, to construct the matrix ***C**^v^* of size *N_s_* × *N_s_*, such that the subjects within a community are assigned the same label.

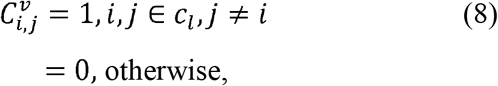

*i, j* = 1, …, *N_S_* and 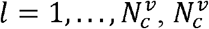 is the number of communities within variable *v*.
2. The multivariate common community is defined as the number of variables in which a pair of participants share a common community. The multivariate common community adjacency matrix of size *N_s_* × *N_s_* is defined as

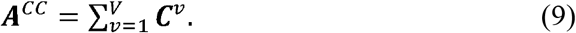
3. The group-wise common community-based affinity for the *p*th participant is calculated as:

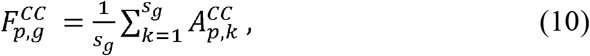

where *g* = 1, …, *G*.

### Implementation on the ASRB data

Data from a total of 195 variables, adjusted for age and sex through generalized linear regression, was used to calculate affinity scores, covering clinical, cognitive, psychosocial and brain structural domains, including cortical and subcortical structures. A complete list of variables and their description is presented in Table 1.

**Table 1:** Metrics of clinical and cognitive assessments.

| **Variable** (*Description*) | **Domain** | **Test** |
| --- | --- | --- |
| LNS (*Working memory*) | Cognition | WMS |
| Attention | Cognition | RBANS |
| Constructional | Cognition | RBANS |
| Delayed memory | Cognition | RBANS |
| Immediate memory | Cognition | RBANS |
| Language | Cognition | RBANS |
| Total | Cognition | RBANS |
| Total scale | Cognition | RBANS |
| Matrix reasoning | Cognition | WASI |
| Vocabulary | Cognition | WASI |
| *Premorbid IQ* | Cognition | WTAR |
| *Verbal fluency* | Cognition | COWAT |
| *Global functioning* | Clinical | GAF |
| Cognitive perceptual | Clinical | SPQ |
| Interpersonal | Clinical | SPQ |
| Disorganized | Clinical | SPQ |
| Social functioning | Clinical | Demographics |
| Social task | Clinical | Demographics |
| LNS, Letter Number Sequencing; WMS, Wechsler Memory Scale; RBANS, Repeatable Battery for Assessment of Neuropsychological Status; WASI, Wechsler Abbreviated Scale of Intelligence; WTAR, Wechsler Test of Adult Reading; COWAT, Controlled Oral Word Association Test; GAF, Global Assessment of Functioning; SPQ, Schizotypal Personality Questionnaire |  |  |

### Diagnostic Verification

The diagnostic verification algorithm was applied to the whole dataset to calculate variable-wise affinity scores using Eqs. 1-2, with each participant included in its neighbourhood. HCs, Scz and TRS individuals were assigned a unique group label. Two diagnostic verification frameworks were developed: a two-group framework (HCs and Scz) and a three-group framework using all three groups. Multivariate metrics of diagnostic accuracy were calculated as described in Eqs. 3-10 to assign affinity-based labels to each individual in the dataset. An individual’s diagnosis was considered verified if the original and affinity-based labels were identical, and atypical if the affinity-based label differed from the original label.

### Classification

To perform affinity-based classification, individuals with TRS were relabeled as Scz, leading to binary classification with groups: HC (n=136) and Scz (n=216). The relabeled data was split into training (110 HCs, 170 Scz) and test (26 HCs, 46 Scz) sets. 5-fold nested cross-validation was performed on the training set, with hyper-parameter tuning performed in the inner loop on the scaling parameters (Eq. 1), using grid search: *α* ∈ {1, …, 10}. In each inner iteration, training data were used to calculate affinity scores, and multivariate affinity-based metrics were calculated (composite affinity, common neighbourhood-based and common community-based metrics). In the validation step, each participant’s affinity-based measures were calculated with reference to the training data, and common neighbourhood-based affinity was used to determine classification accuracy: the number of correctly labelled participants in the validation set divided by the total sample size. The outer loop was used to calculate average classification accuracy across folds.

For comparison, weighted K-nearest neighbours (KNN) classification was performed using MATLAB’s *fitcknn* function, and nested cross-validated on the same partitions of the training data. For weighted KNN classification, weighting was performed using the ‘*inverse distance*’ option on the standardized Euclidean distance between a participant and its *k*th neighbour, and hyper-parameter tuning was performed on the number of neighbours, *K*, using grid search: *K* ∈ {5, …, 15}. A schema of the nested cross-validation algorithm is presented in Supplementary Figure S1.

The affinity-based and weighted KNN models were tuned again on the entire training dataset to set hyperparameters, instead of selecting the best-tuned model from the nested cross-validation. As both affinity-based and weighted KNN classifiers are sensitive to the local neighbourhood composition, changes in sample-size require re-tuning of hyperparameters to ensure optimal performance. *α* = 6 and *K* = 6 showed highest accuracy on the training set. The tuned models were applied to the independent test set and prediction accuracy of both classifiers was calculated.

### Post-hoc analyses

To assess the classification separability of the TRS participants, the original group labels were re-applied post-hoc to the multivariate affinity scores from the training data. K-means clustering was performed in MATLAB^26^, and three separate clusters were identified using multivariate composite affinity, common-neighbourhood and common-community scores as features. The within cluster group composition was assessed to determine the separability of TRS individuals from other groups.

## Results

### Diagnostic Verification

In the two-group diagnostic verification framework, the HCs had verified diagnostic status more frequently when compared to the chronic schizophrenia participants (Table 2). The composite multivariate affinity metric was most sensitive to the heterogeneity within the Scz group, with 64.83% of individuals receiving a verified diagnosis. The vote-based affinity metric was least sensitive to within group heterogeneity, leading to ~100% verified diagnoses.

**Table 2:** Performance evaluation of affinity-based diagnostic verification framework across two diagnostic groups

| Metric | Diagnostic verification (%) |  |  |  |
| --- | --- | --- | --- | --- |
|  | HC |  | Scz |  |
|  | Verified | Atypical | Verified | Atypical |
| Composite multivariate affinity | 97.05 | 02.95 | 64.83 | 35.17 |
| Common Neighbourhood-based affinity | 94.85 | 05.15 | 73.07 | 26.93 |
| Common Community-based affinity | 92.64 | 07.36 | 74.17 | 25.83 |
| Vote-based multivariate affinity | **100.00** | **0.00** | **100.00** | **0.00** |

In the three-group diagnostic verification framework, the HCs and TRS participants had the most frequently verified group membership (Table 3). Similar to the two-group framework, the Scz participants were least likely to achieve a verified diagnosis, potentially due to a higher degree of overlap with both HC and TRS groups and imbalanced sample-sizes, except when the vote-based affinity metric was used (99.5% diagnostic verification). The common community-based affinity metric was the strictest metric, with only 39.56 % verified Scz participants.

**Table 3:** Performance evaluation of affinity-based diagnostic verification framework across three diagnostic groups

| Metric | Diagnostic verification (%) |  |  |  |  |  |
| --- | --- | --- | --- | --- | --- | --- |
|  | HC |  | Scz |  | TRS |  |
|  | Verified | Atypical | Verified | Atypical | Verified | Atypical |
| Composite multivariate affinity | 97.79 | 02.21 | 65.93 | 34.07 | 97.05 | 02.95 |
| Common Neighbourhood-based affinity | 92.64 | 07.36 | 43.40 | 56.60 | 67.64 | 32.36 |
| Common Community-based affinity | 89.70 | 10.30 | 39.56 | 60.44 | 91.17 | 08.83 |
| Vote-based multivariate affinity | **100.00** | **0.00** | **99.45** | **0.55** | **100.00** | **0.00** |

**Table 4:** Classification accuracy of affinity-based and weighted KNN classifiers in the training, nested cross-validated and prediction datasets. Common neighbourhood and community-based classification outperform composite multivariate affinity and weighted KNN classification.

| Classifier | Metric | Accuracy (%) |  |  |  |  |  |  |  |  |
| --- | --- | --- | --- | --- | --- | --- | --- | --- | --- | --- |
|  |  | Training |  |  | Nested Cross-validation |  |  | Prediction |  |  |
|  |  | HC | Scz | Total | HC | Scz | Total | HC | Scz | Total |
| Affinity-based Classifier | Composite multivariate affinity | 95.45 | 58.82 | 73.21 | 90.91 | 70.59 | 78.57 | 100.00 | 60.87 | 75.00 |
|  | Common Neighbourhood-based affinity | 94.55 | 71.76 | 80.71 | 91.82 | 75.29 | 81.79 | 96.15 | 78.26 | 84.72 |
|  | Common Community-based affinity | 95.45 | 71.18 | 80.71 | 88.18 | 74.12 | 79.64 | 96.15 | 82.61 | 87.50 |
| K-Nearest Neighbours | Standardized Euclidean distance | 100.00 | 100.00 | 100.00 | 82.73 | 77.65 | 77.86 | 80.77 | 76.09 | 77.78 |

**Table 5:**
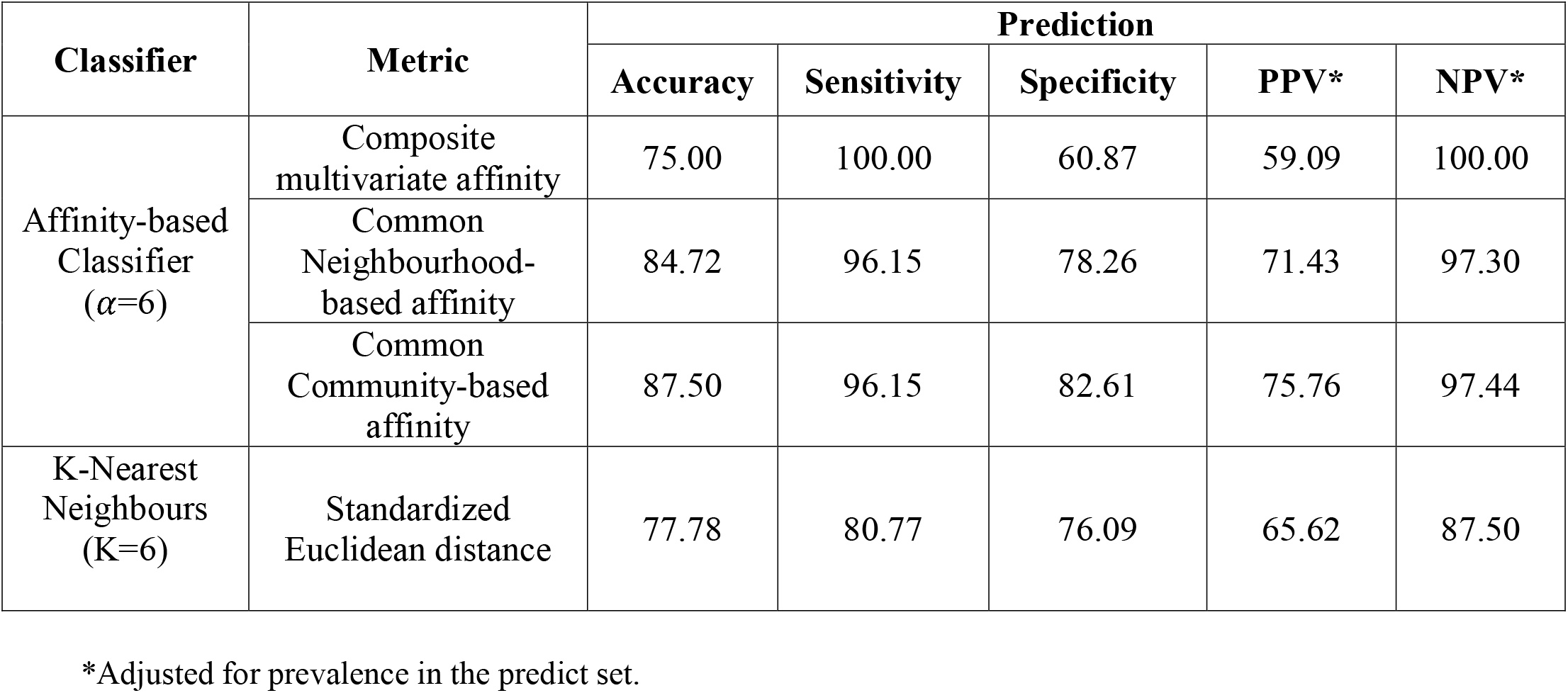
Performance evaluation of affinity-based and weighted KNN classification in the prediction dataset

| Classifier | Metric | Prediction |  |  |  |  |
| --- | --- | --- | --- | --- | --- | --- |
|  |  | Accuracy | Sensitivity | Specificity | PPV* | NPV* |
| Affinity-based Classifier ($\alpha=6$) | Composite multivariate affinity | 75.00 | 100.00 | 60.87 | 59.09 | 100.00 |
|  | Common Neighbourhood-based affinity | 84.72 | 96.15 | 78.26 | 71.43 | 97.30 |
|  | Common Community-based affinity | 87.50 | 96.15 | 82.61 | 75.76 | 97.44 |
| K-Nearest Neighbours (K=6) | Standardized Euclidean distance | 77.78 | 80.77 | 76.09 | 65.62 | 87.50 |

\*Adjusted for prevalence in the predict set.

For each individual, the variable-wise affinity scores form a multidomain ‘*fingerprint*’ demonstrating within variable affinity of that individual to each diagnostic group in the data (Figure 1). The exemplar affinity fingerprint from a healthy participant explains the individualized attributes, with predominantly healthy structural, cognitive and clinical affinities. In contrast, the exemplar fingerprints from chronic schizophrenia and TRS individuals reveal a more heterogeneous pattern of group affinities, providing the evidence for an individual-centric multi-domain multivariate pattern capable of taking intra-illness heterogeneity and inter-illness overlap into account.

**Figure 1:**
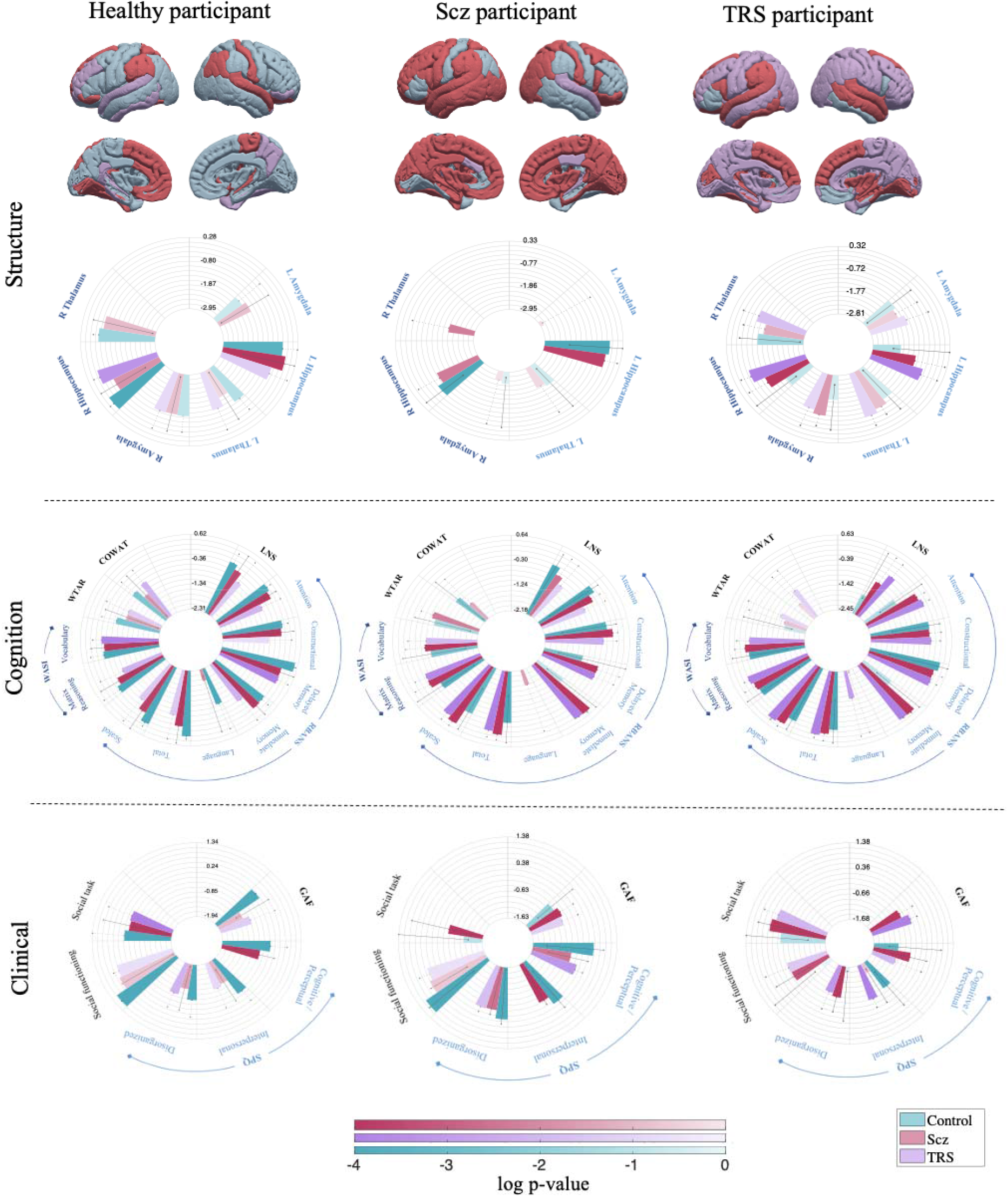
Exemplar individual-centric affinity fingerprints. The cortical pattern describes maximal group affinity within each region. In the circular bar graphs, the variable-wise subcortical, cognitive and clinical affinity scores are plotted on the log-scale with 95% confidence intervals (absent bar: zero affinity score, dotted errorbar: zero lower CI). Shading on the bars is based on the log p-value quantifying score reliability. Within each variable, the largest bar corresponds to the group to which the individual shows strongest affinity, followed by the group with second-highest affinity and so on.

### Classification

The affinity scores-based classification achieved a high degree of accuracy in the training, nested cross-validation and prediction steps (Table 2). The common neighbourhood (cross-validated accuracy: 81.79%, prediction accuracy: 84.72%) and common community based (cross-validated accuracy: 79.64%, prediction accuracy: 87.50%) metrics outperformed the weighted KNN algorithm (cross-validated accuracy: 77.86%, prediction accuracy: 77.78%), with the exception of nested cross-validated accuracy in the Scz samples where the weighted KNN classifier achieved slightly higher accuracy (77.65%) than the common-neighbour (75.29%) and common-community (74.12%) affinity-based classification. In the prediction dataset, affinity-based classification achieved higher sensitivity (common-neighbour: 96.15%; common-community: 96.15%) and specificity (common-neighbour: 78.26%; common-community: 82.61%) and more robust positive predictive value (PPV; common-neighbour: 71.43; common-community: 75.76) and negative predictive value (NPV; common-neighbour: 97.30; common-community: 97.44), when accounted for group prevalence in the prediction sample (Table 3).

### Separability of TRS participants

In the classification step, the separability of TRS participants from the healthy controls and chronic schizophrenia participants was assessed post-hoc by relabeling the TRS participants from Scz to TRS in the training data (Figure 2). On average, TRS participants showed low affinity to both HCs and Scz participants (Figure 2 A-B), suggesting that TRS participants are separable from these groups. To establish the degree of separability of the TRS participants, k-means clustering was performed on the training dataset using multivariate composite affinity, common-neighbourhood and common-community scores. Three clusters were identified, with cluster 1 predominantly occupied by Scz participants, cluster 2 by TRS participants and cluster 3 by HCs (Figure 2 C-D).

**Figure 2:**
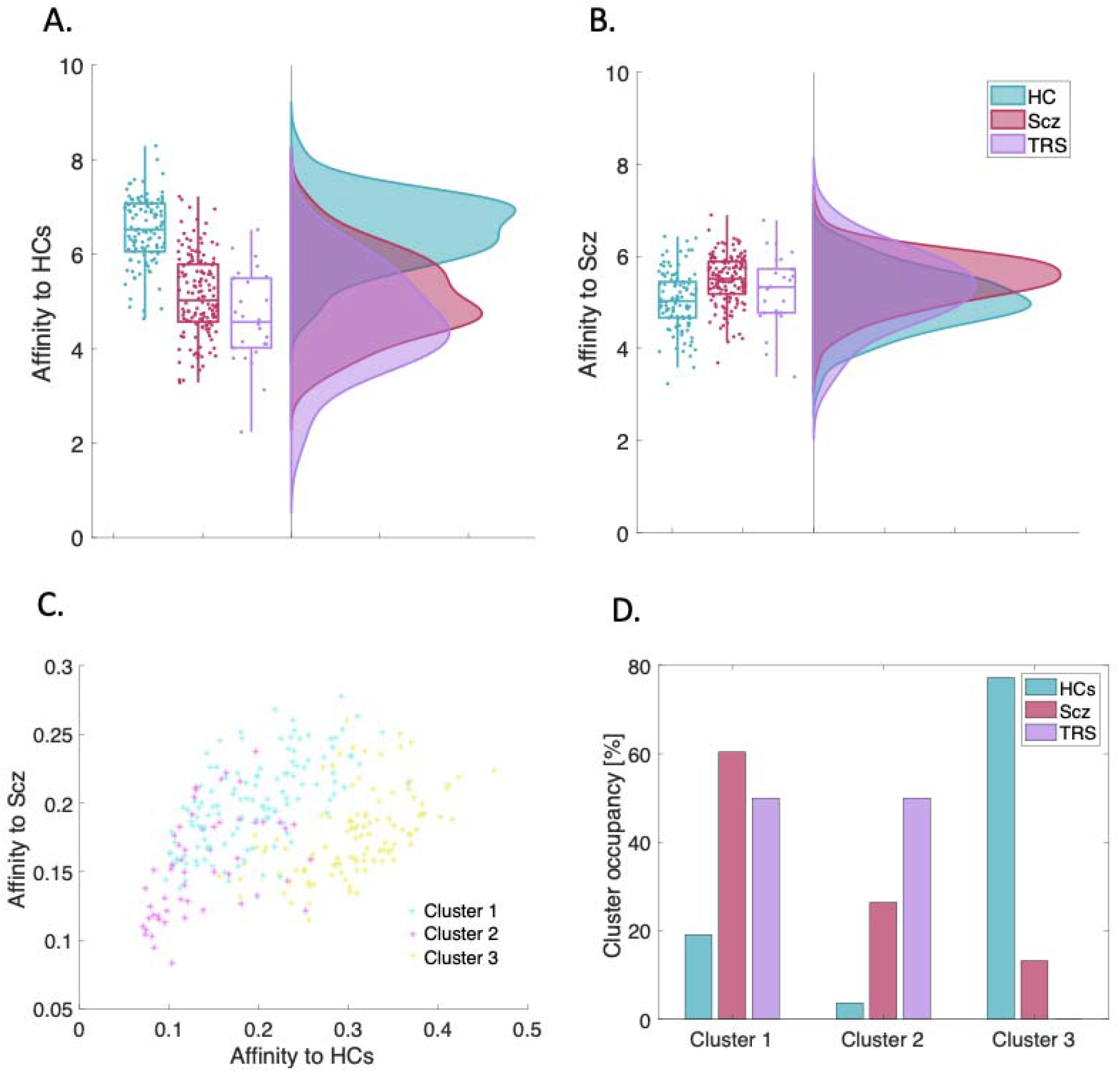
Classification separability of TRS participants from healthy controls and Scz participants in the training dataset (metric: common neighbourhood-based affinity). TRS individuals showed low affinity to both (A) HCs and (B) Scz groups. C) Post-hoc K-means clustering on the multivariate scores from the training dataset used for classification. Three clusters were identified based on affinity to HC and Scz groups using scores from composite affinity, common neighbourhood and common community-based metrics. D) Group-wise composition of the three clusters (number of participants per group divided by group sample size in the training data).

## Discussion

In this study, we introduce the affinity score framework and examine its utility for accurate verification and classification of schizophrenia-spectrum disorders. An individual’s affinity profile provides a snapshot of their biopsychosocial marker affinity to different diagnostic groups across a wide range of measures, including clinical, cognitive, social, and neuroimaging domains. Furthermore, the classification accuracy of both our verification and classification algorithms was high. Specifically, our verification algorithm was successfully able to verify ~100% of study participants using the voting-based affinity metric, and our classification algorithm successfully classified 91% of healthy controls and 72% of schizophrenia participants.

Traditional mean-centric approaches are based on assumptions of mutual exclusivity between diagnoses and intra-illness homogeneity and are therefore incapable of taking into account individual-level variation across biopsychosocial measures. Our affinity score framework, on the other hand, takes an individual-centric approach to understanding psychiatric disorders, and is therefore capable of accounting for factors such as comorbidity, and intra-and inter-group heterogeneity and overlap. Across a range of measures, each individual is considered to be the centre of their own unique neighbourhood, with no assumption that the composition of these neighbourhoods will be the same across measures. Thus, one individual may have high affinity to individuals with TRS on one measure, and high affinity to healthy individuals on another. Such an approach allows clinicians to identify not only the most salient treatment targets for any given individual, but also the areas of strength that might be leveraged to enhance treatment efficacy. This approach also allows clinicians to visualise at a glance the diagnostic group/s to which an individual is most closely aligned across multiple variables, aiding in the verification or confirmation of diagnoses and potential comorbidities.

In two-group analyses, our diagnostic verification approach, which incorporates cognitive, clinical, functioning, and neuroimaging measures, showed very high capability for the verification of schizophrenia and healthy control groupings using the voting-based metric (100% for both groups). However, other metrics, including common neighbourhood- and common community-based affinity were more sensitive to within group heterogeneity, particularly for individuals with schizophrenia, resulting in lower verification capability (73.07% and 74.17% respectively). Similar findings were observed for the three-group analysis, which included TRS individuals. Here, the vote-based metric again showed very high diagnostic verification capability across all three groups. As with the two-group analysis, verification capability was lowest for individuals with schizophrenia, particularly for common neighbourhood- and common community-based metrics, suggesting a greater degree of heterogeneity among individuals with schizophrenia compared to healthy controls or individuals with TRS. Together, these findings suggest that the composite affinity, common neighbourhood- and common community-based frameworks might be useful for identifying intra-diagnosis heterogeneity within the schizophrenia subgroup, where a proportion of patients more closely resemble healthy individuals when the strength of their affinity scores are taken into account. The vote-based metric, on the other hand, while less sensitive to intra-diagnosis heterogeneity, may be particularly useful for diagnostic verification. Together, these metrics would serve as particularly useful clinical decision support tools as they would allow clinicians to determine the overlap between their proposed diagnosis for an individual and the likely diagnosis or diagnoses determined by that individual’s affinity scores. In cases where there is a discrepancy between the clinician’s diagnosis and the diagnosis provided by the verification algorithm, the clinician would be able to determine whether the individual is an atypical presentation of their proposed diagnosis, or whether there may be comorbidities or a differential diagnosis relevant to the individual.

Using all the available biopsychosocial measures, the accuracy of our classification algorithm was also very high. Due to the sample size requirements of classification algorithms, the small number of TRS individuals included in this study were merged with the larger schizophrenia group for the purposes of classification. Unlike the verification algorithm, in which individuals are considered part of their own neighbourhoods, classification requires that individuals are not counted towards their affinity scores, and they are therefore not included in their neighbour counts. In our test sample of 66 schizophrenia participants and 26 healthy controls, our overall classification accuracy was 87.5% for common community-based affinity and 84.72% for common neighbourhood-based affinity, which can be compared to 77.78% for the KNN algorithm. The high level of accuracy observed using our classification algorithm suggests this framework has potential as a diagnostic prediction tool in a clinical setting. Of note, when TRS individuals were re-labeled as TRS following affinity score calculation, they did not have high affinity to either the healthy control group or the schizophrenia group. Thus, our algorithm demonstrates the capability of identifying disease subgroups or under-represented populations.

Due to a high degree of conceptual and algorithmic overlap, we compared the performance of affinity-based classification with the weighted KNN classification algorithm. Both algorithms are sensitive to localised information in the data, in contrast to other popular classifiers such as support vector machines, which attempt to identify separation boundaries between classes^27^. Further, prior to classification, no feature selection was performed, and all available variables were included in the diagnostic verification and classification algorithms. Although a more efficient feature selection strategy can potentially lead to improved classification accuracy, less discriminant features still contain clinically-relevant information and excluding such features contradicts the aim of developing a clinically-focused diagnostic support system based on the affinity scores.

In a final step, k-means clustering was applied to multivariate affinity score profiles in order to assess the separability of treatment-resistant individuals from healthy controls and individuals with chronic schizophrenia. The three clusters derived from individual biopsychosocial Affinity profiles included a largely healthy cluster, and two clusters dominated by the schizophrenia and TRS groups. The first of these clusters appears to represent moderate illness severity: the largest contribution to this cluster comes from individuals with schizophrenia, although there is also a relatively large contribution from the TRS group. Without adequate symptom measures it is difficult to fully characterise this group, however, it is possible that the TRS individuals in this cluster are those who have responded to clozapine treatment and are thus functioning at a higher level than those who do not respond to clozapine. The final cluster is largely made up of TRS individuals, with a smaller group of individuals with schizophrenia also present. This cluster most likely represents the most severe illness phenotype, and the individuals with schizophrenia within this cluster may be missed TRS individuals due to the use of clozapine prescription as our grouping measure, or simply a more severely affected subgroup of individuals with treatment-responsive schizophrenia. Thus, as with our verification and classification frameworks, this clustering analysis highlights that, while healthy individuals tend to cluster together, heterogeneity in biopsychosocial profiles exists across the schizophrenia-spectrum.

The affinity score framework has several benefits over more traditional population-based approaches. One important benefit is that this framework is entirely data-driven and does not require individual scores to fall into any particular type of distribution - there is no assumption of “normality”. Furthermore, unlike traditional statistical frameworks, outliers will not have a negative impact on the affinity score algorithm and are considered to be data points of interest rather than scores to be removed or adjusted. Affinity scores can also be dynamic, both at the individual and the population level. At the individual level, affinity scores and ‘*fingerprints*’ can change over time as an individual’s illness evolves, and could potentially be used to predict illness-related outcomes over time. At the population level, the affinity score framework is capable of dealing with the addition of new diagnostic groups and new individuals within existing diagnostic groups, providing a flexible approach to biomarker identification. Thus, although we have developed the affinity score framework in a sample of individuals with schizophrenia, it could be equally applied to other clinical populations, including adolescents and adults, and would have particular utility in the early stages of psychiatric illness where symptoms are often less specific. There is no limit to the number of diagnoses that could be included in a model. This system therefore represents a significant breakthrough in knowledge and practice, substantially improving diagnosis and treatment of psychiatric disorders across diagnostic boundaries.

Our findings should be considered in the context of several limitations. In this sample, TRS was defined based on clozapine prescription, however it is possible that some individuals with schizophrenia who are actually treatment-resistant may not have been prescribed clozapine, and therefore would have incorrectly been included the schizophrenia group. Furthermore, given the small sample size of the TRS group, it was not possible to include these individuals as a separate diagnostic group for the purpose of diagnostic classification and they were therefore merged with the schizophrenia group for this step. However, given that affinity scores are calculated in standardised space, it will be possible to merge in new datasets for future testing and development of the algorithm. In addition, recent developments in harmonization techniques for neuroimaging data, such as the ComBat pipeline^28,29^, will allow for the inclusion of MRI data acquired on different scanners. It is also possible to construct partial fingerprints using only the available measures for a single individual, ensuring missing data does not preclude someone from being included in the Affinity Framework.

The historical categorical, diagnostic approach contrasts with the hierarchical taxometric dimensional method in the description of psychopathology and mental illness^30^. Both systems, however, are based on the grouping of individual symptoms and characteristics, constraining them alternatively to a defined disorder, or to a statistical pattern of characteristics, hierarchically shared by all. Both, consign the individual to a collective. They fail to adequately capture the individual nature of either their comorbidity and heterogeneity or their specific biopsychosocial aetiology, phenomenology, treatment specificity, and developmental pathway. The individual is lost in the collective of both the categorical and dimensional attempts to understand psychopathology. Individual affinity “fingerprinting” aims to bring an individual focus to the categorical and dimensional approaches to understanding psychopathology and thus inform research and clinical practice.

## Author Contribution Statements

Cassandra Wannan: Data curation, conceptualization, writing - original draft preparation, review and editing.

Antonia Merritt: Project administration, data collection, review and editing.

Christos Pantelis: Supervision, conceptualization, funding acquisition, review and editing.

Warda Syeda: Conceptualization, formal analyses, methodology and algorithm development, software implementation, visualization, writing - original draft preparation, review and editing.

## Acknowledgements

This study was supported by the Australian Schizophrenia Research Bank (ASRB), Chief Investigators: Carr VJ, Schall U, Scott R, Jablensky A, Mowry B, Michie P, Catts S, Henskens F, Pantelis C, Loughland C. The ASRB is supported by Neuroscience Research Australia (NeuRA). CP was supported by a National Health and Medical Research Council (NHMRC) L3 Investigator Grant (1196508), and NHMRC Program Grant (ID: 1150083).

## Supplementary Material

### Methods

#### Sample-size requirements of affinity-based classification

Sample size calculations of the affinity-based classification were performed using the method described in (Figueroa et al. 2012).

#### Permutation testing

To assess the interchangeability of variable-wise affinity scores between groups, permutation testing was performed with the null hypothesis that affinity scores are interchangeable between groups. An iterative algorithm was implemented:

1. For each iteration, the group labels of all subjects were randomly permuted using MATLAB’s *randperm* function.
2. Variable-wise affinity scores were calculated, as described in the Methods section.
3. Steps 1-2 were repeated 10,000 times, with each iteration having randomly permuted group labels.
4. A p-value for the *p*th subject, *g*th group and *v*th variable was calculated based on the total number of times the permuted affinity score was greater than or equal to the estimated affinity score:

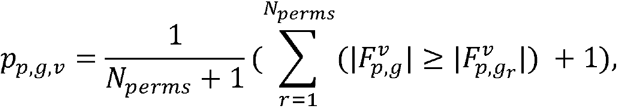

where *g* is the original group label and *g_r_* is the permuted group label.

#### Bootstrapping

Bootstrapping was performed to estimate confidence intervals of the variable-wise affinity scores, using an iterative algorithm:

1. In each iteration, random sampling with replacement was performed to generate samples from the original data from all variables. The number of samples per group was the same as the original data.
2. For each bootstrap sample, the variable-wise affinity scores were calculated, and the process was repeated 10,000 times to generate a sampling distribution of the variable-wise affinity scores.
3. The 95% confidence intervals were calculated from the sampling distribution.

#### Nested Cross-validation

**Figure S1:**
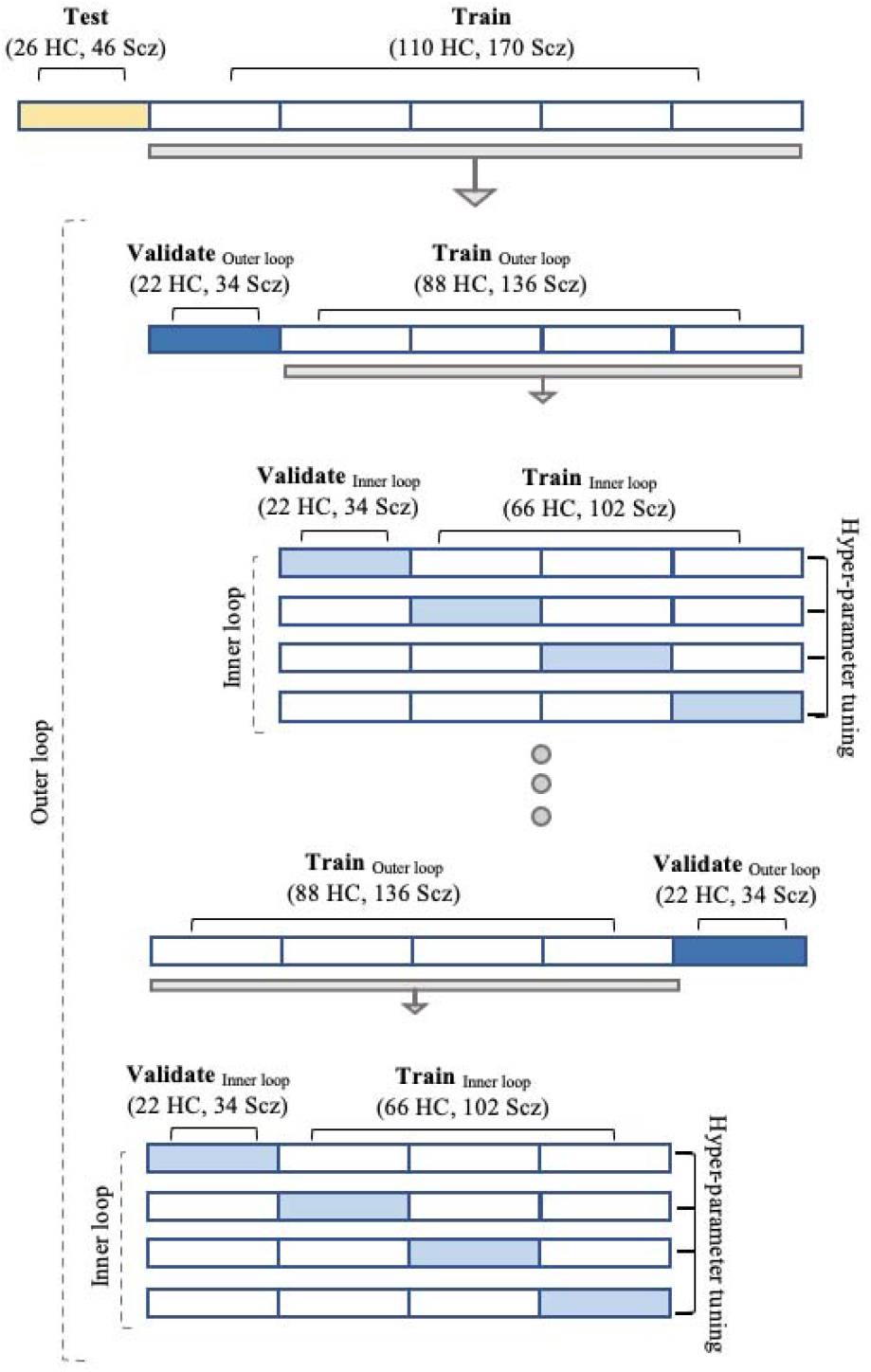
Nested cross-validation. 5-fold nested cross-validation was performed on the training set, with hyper-parameter tuning performed in the inner loop on the scaling parameters, and K for the affinity-based and K-nearest neighbours algorithms, respectively. The outer loop was used to calculate average classification accuracy across folds.

